# Reevaluating the Role of Beta2-Microglobulin: New Insights on Selective Vulnerability in ALS Pathology

**DOI:** 10.1101/2025.11.27.690988

**Authors:** M Leboeuf, J Nijssen, LH Comley, JC Aguila Benitez, I Mei, S Gómez Alcalde, V Radoi, S Nichterwitz, C Schweingruber, E Hedlund, S Cullheim

## Abstract

Amyotrophic lateral sclerosis (ALS) is characterized by the selective loss of motor neurons (MNs). Why these neurons show particular vulnerability in ALS is still debated, and also why certain MNs remain resilient. We investigated the importance of the human leukocyte antigen (HLA) and particularly beta2-microglobulin (β2m) in MN susceptibility to ALS as these molecules have been implicated both in prolonging and shortening disease. RNA-sequencing demonstrated that disease-resistant oculomotor neurons (OMNs) and Onuf’s MNs had similar *β2m* and *HLA* mRNA levels to vulnerable spinal MNs in control tissues. Thus, baseline differences in these transcripts do not explain the differential vulnerabilities of these neurons to disease. Nonetheless, the HLA protein level showed an inverse correlation with spinal MN size, where large MNs, which are lost early in ALS, had low HLA levels. HLA protein levels were also reduced in end-stage ALS patient spinal MNs, while remaining unchanged in OMNs with disease. Spinal MNs uniquely exhibited a significant upregulation of *β2m* and *HLA-C* transcripts with disease, likely as a protective compensatory response. Thus, β2m and HLA may relate to vulnerability to ALS. Cross-breeding of β2m knockout mice with SOD1G93A ALS mice resulted in modest transient worsening of functional motor performance, but did not affect life span. There was partial preservation of innervation of particular muscles in SOD1G93A mice lacking β2m, which was insufficient to improve motor behavior. These findings indicate that β2m and HLAs are dynamically regulated in ALS, and may influence MN vulnerability, but they are not major disease modifiers in ALS.

## Introduction

Major histocompatibility complex class I (MHC-I) molecules are present as transmembrane glycoproteins on the surface of all nucleated cells. MHC-I proteins display peptide fragments to circulating cytotoxic T cells and natural killer cells (NKs). This antigen presentation is essential for immune defense against intracellular pathogens and cancer (Neefjes et al. 2011). In case the peptide fragments derive from non-self-proteins, this may trigger an immediate detrimental response from the immune system. However, alterations in MHC-I expression can have unintended consequences; downregulation can trigger NK cell-mediated cytotoxicity (Höglund et al. 1997), whereas aberrant upregulation is implicated in autoimmunity and neurodegeneration (Cebrián et al. 2014). Structurally, MHC-I consists of an alpha heavy chain, which has two domains to which peptides bind, an immunoglobulin (Ig)-like domain and a transmembrane region with a cytoplasmic terminus. The heavy chain of class I molecules is encoded by genes at the HLA-A, HLA-B or HLA-C loci in humans, and is linked to a conserved light chain, beta-2-microglobulin (β2m), which is required for MHC-I stability and surface expression (Zijlstra et al. 1990) (Figure 1A). Beyond its classical immunological role, MHC-I has been increasingly recognized as a key player in the nervous system, where it contributes to synaptic pruning, neurodevelopment, plasticity (Boulanger 2004; Corriveau et al. 1998; Huh et al. 2000; Goddard et al. 2007; Shatz 2009; Oliveira et al. 2004) and a possible involvement in neurological disorders (Haines et al. 1998; Kim et al. 2023; Afify et al. 2024). Motor neurons, in particular, exhibit high expression of mRNAs for MHC-I, including β2m, under physiological conditions (Lindå et al. 1998), with further upregulation following axonal injury where MHC-I protein accumulates at the end of the cut axons (Lindå et al. 1998; Thams et al. 2009). The mouse gene H2-Db also increases in motor neuron (MN) somas after axotomy (both at mRNA and protein level) and in axons (Thams et al. 2009). These findings suggest that MHC-I may influence neuronal survival and degeneration. The presence of high levels of MHC-I and β2m in MNs has spurred an interest in finding a role for these proteins in degenerative MN diseases, in particular amyotrophic lateral sclerosis (ALS). While MHC-I and β2m are suspected to play a role in ALS, research has yielded diverse findings regarding their expression and function. In an ALS mouse model, expressing a disease-associated variant of superoxide dismutase 1 (SOD1G93A) resulted in increased *β2m* mRNA levels in spinal MNs of symptomatic mice. When the SOD1G93A mice were crossed with β2m knockout mice, survival was decreased due to a shorter disease duration, indicating that β2m was protective in ALS and that upregulation of β2m on MNs in ALS may be a protective compensatory response. However, this study did not investigate the level of MHC-I or β2m on the protein level (Staats et al. 2013). Another study reported that HLAs were reduced in MNs in symptomatic SOD1G93A mice and that MHC-I levels were completely abolished in MNs in end-stage ALS patient tissue (Song et al. 2016). It was moreover demonstrated that astrocytes reduced HLA expression on MNs, rendering them more susceptible to astrocyte-induced cell death. Overexpression of HLA-C (H2-K1 in mouse) in MNs, but not HLA-A or HLA-B, could rescue this susceptibility (Song et al. 2016). The abovementioned studies suggest that the presence of β2m or HLAs, and presumably then MHC-I, on MNs may have a beneficial effect on MN survival in ALS. On the other hand, a third study suggested that while the activation of MHC-I in the peripheral nervous system of SOD1G93A mice preserves muscle innervation and motor function, the effects in the central nervous system (CNS), mediated by the interaction of microglia expressing MHC-I with CD8^+^ T cells accelerates MN death and reduce the overall survival of SOD1G93A mice and thus removing β2m increased survival of the mice (Nardo et al. 2018). These disparities highlight the complexity of MHC-I regulation in ALS and underscore the need for further investigation.

**Figure 1.**
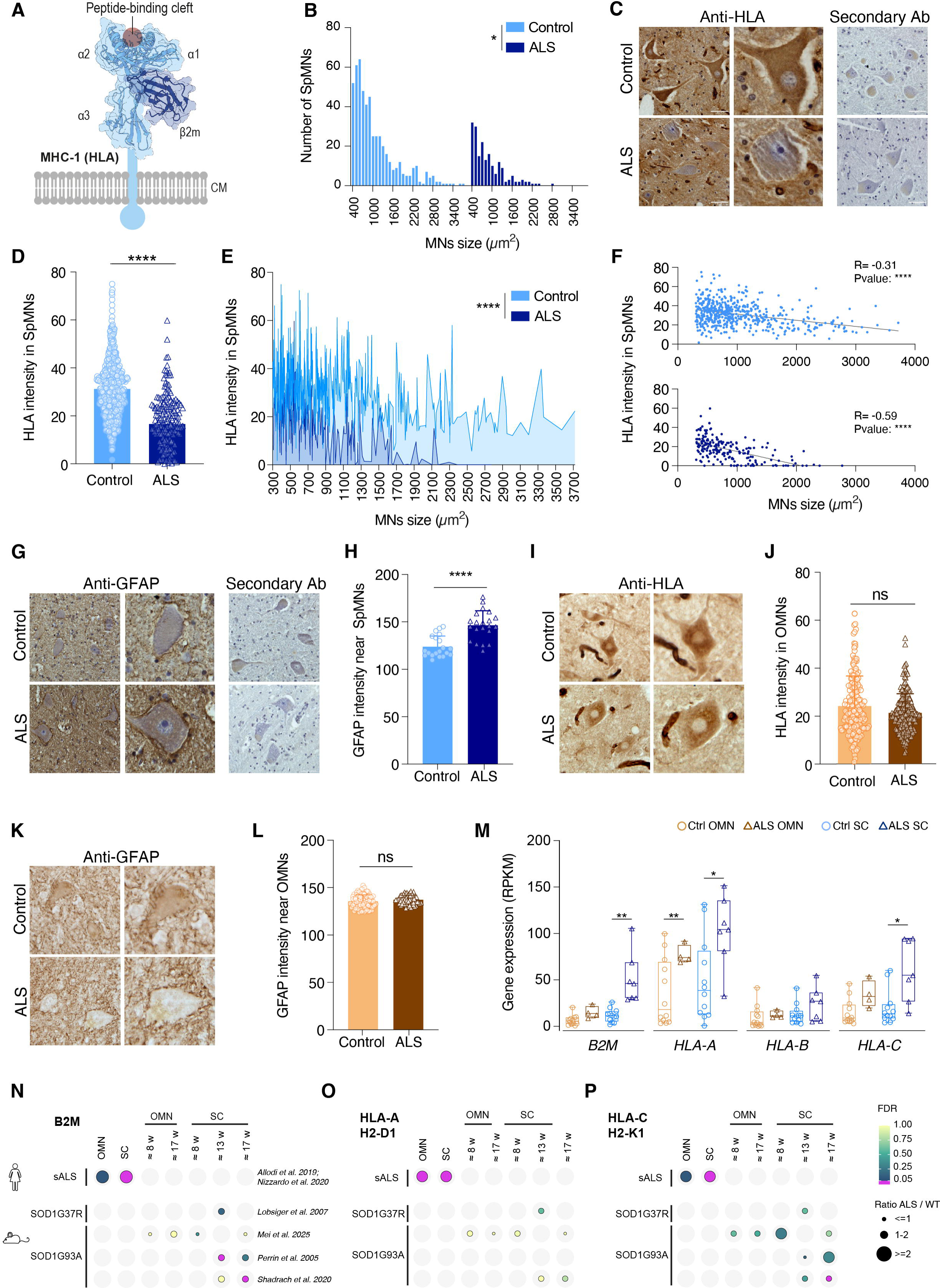
Motor neuron loss in ALS is correlated with a decrease in HLA protein, increased GFAP immunoreactivity and upregulation of *B2M* and HLA-A/C mRNAs. (**A**) Schematic of the MHC molecule. (**B**) Measurement of motor neuron area (μm2) in control and ALS spinal cord donor tissues shows a loss of motor neurons across sizes (*P<0.05,* Kolmogorov-Smirnov test). (**C**) Representative images of immunohistochemical staining against HLA in human control and ALS spinal cord *post mortem* donor tissues. (**D**) Quantification of HLA immunoreactivity from antibody-staining (in C) shows a significant decrease in ALS end-stage patient spinal motor neurons (SpMNs) compared to control (P*<0.0001,* Mann-Whitney Unpaired t-test, data are expressed as the mean ± SD). **(E**) HLA intensity in spinal motor neurons correlated with cell size (*P<0.0001*, Kolmogorov-Smirnov Unpaired t-test). (**F**) HLA intensity and spinal motor neuron size are inversely correlated both in human control and ALS spinal cord *post mortem* donor tissues (Ctrl: R=-0.35, *P<0.0001*, Spearman correlation, ALS: R=-0.57, *P<0.0001*, Spearman correlation) (**G**) Representative immunohistochemical staining against GFAP in human control and ALS spinal cord donor tissues. (**H**) Quantification of GFAP immunoreactivity in the spinal cord shows an increase in ALS patients (*P<0.0001,* Mann-Whitney Unpaired t-test, data are expressed as the mean ± SD). (**I**) Representative images of immunohistochemical staining against HLA in human control and ALS brain donor tissues. (**J**) Quantification of HLA immunoreactivity from antibody-staining (in I) shows no difference in HLA intensity in the oculomotor neurons (OMNs) between control and ALS donors (*P=0.06,* Mann-Whitney Unpaired t-test, data are expressed as the mean ± SD). (**K**) Representative images of immunohistochemical staining against GFAP in human control and ALS donor tissues in the OMNs region. (**L**) Quantification of GFAP immunoreactivity shows no difference between control and ALS donors in the OMNs region (*P=0.11,* Mann-Whitney Unpaired t-test, data are expressed as the mean ± SD). (**M**) RNA sequencing of MNs shows an increase in *β2M* levels in spinal MNs in ALS, and an increase in *HLA-A* in OMNs and spinal MNs with disease as well as an increase in *HLA-C* in spinal MNs in ALS. (N = min. 4 samples per group, each representing one individual. P values: ** < 0.01, *** < 0.001, obtained with 2-way ANOVA). (**N-P**) Summary of (**N**) *β2M*, (**O**) *HLA-A/H2-D1* and (**P**) *HLA-C/H2-K1* expression levels in OMN and spinal MNs (SC) across sporadic ALS (sALS) and mutant SOD1 ALS mice.

To address this, we analyzed MHC-I expression in ALS across human and mouse datasets. Our analysis shows that there is an inverse correlation with HLA protein levels and MN size in the spinal cord. We also show that HLA protein levels are reduced in spinal MNs with ALS and that this is unique to vulnerable MNs while OMNs maintain their expression throughout disease. We also demonstrate a compensatory upregulation of *β2m* and *HLA-C* mRNAs only in ALS-vulnerable spinal MN, which may be part of an insufficient neuroprotective response. To directly assess the impact of β2m on ALS disease progression, we generated SOD1G93Axβ2m-/- mice and evaluated their motor function, disease course, and neuronal survival. Our findings indicate that loss of β2m leads to a small early worsening in motor function both in males and females, but does not affect overall survival in SOD1G93A mice. Additionally, we observed that while neuromuscular junction (NMJ) innervation remained unchanged in tibialis anterior and soleus muscles, the lumbrical muscles of SOD1G93Axβ2m-/- mice exhibited increased NMJ stability, suggesting a muscle-specific protective effect, which did not give a functional outcome in our assays.

## Methods

### Bioinformatics analysis of RNA sequencing and microarray data

Processed files of microarray datasets from (GSE40438, Brockington et al. 2013; GSE52118, Kaplan et al. 2014; Perrin et al. 2005; Lobsiger et al. 2007), and processed data sets from Shadrach et al. 2021; Allodi et al. 2019; Nizzardo et al. 2020 were retrieved from the authors. Raw datasets from (Bandyopadhyay et al. 2013; Mei et al. 2025) were mapped to mm39 genome assembly using STAR (version 2.7.0e) (Dobin et al. 2013). Expression levels were determined using the rpkmforgenes.py software with the Ensembl gene annotation (mm39).

### Ethics statement and animal models

All animal procedures were approved by the Swedish ethical council (Stockholms Norra Djurförsöksetiska nämnd) in compliance with US National Institutes of Health guidelines. Mice were housed according to standard conditions, with access to food and water ad libitum and a dark/light cycle of 12 hours at the laboratory animal facility, Karolinska Institutet. To investigate the role of beta 2 microglobulin (β2m) in ALS we used the strain B6.129P2-β2m tm1/Unc/J, Jackson Laboratory, Stock Number 002087. These mice are homozygous for the *B2m^tm1Unc^* targeted mutation (β2m-KO) and fail to express MHC class I protein on the cell surface and are grossly deficient in CD45-CD8+ T cells (Koller et al. 1990). To generate experimental ALS-mice with different doses of β2m we crossed β2m-/- females with transgenic males of the SOD1^G93A^ colony (B6.Cg□JTg(SOD1*G93A)1Gur/J; Jackson Laboratory Stock, Number 004435). These mice overexpress mutant human SOD1 protein leading to an ALS□Jlike phenotype (Gurney et al. 1994). Litters of this first cross (F1), always heterozygous for β2m (β2m+/-), were crossed to generate all possible genotype combinations with the mutant SOD1 background (F2): SOD1^G93A^;β2m+/+, SOD1^G93A^;β2m+/- and SOD1^G93A^;β2m-/-. Some experimental animals, normally produced at lower frequency, were also generated by crossing: β2m-/- females from the original colony with mutant SOD1-transgenic males from the F1 (SOD1^G93A^;β2m+/-) or β2m+/+ females with males (SOD1^G93A^;β2m+/+), both from the F2.

Genotyping of both colonies and experimental animals was carried out by regular PCR following The Jackson Laboratory protocols while qPCR was routinely used to follow transgenic SOD1 copy number of male breeders (also following Jackson’s recommendations).

### Behavioral analysis

Body weight of experimental animals was measured twice a week from postnatal day (P) 56. From week 8 (∼P56) to week 20 (∼P140), animals were weekly scored for the extension reflex of their hind limbs and challenged with the inverted grid test. In the extension reflex test, the mice were suspended from the tail and their hind limb extension reflex was scored from 3.0 (normal) to 1.0 (ALS condition) with 0.5 interval points during disease progression. In the inverted grid test we measured the time a mouse holds a grid, while suspended over a 40cm high cylinder. Maximum duration was 120 seconds (healthy, normal situation after task learning). If the animal did not reach the 120 seconds, another trial was carried out after 1-2 minutes of recovery. The longest time measured was selected as the behavioral performance for that week. Some records were excluded from the inverted grid analysis since a few animals learn to jump after just a few seconds of trial. Ultimately, maximum performance of each mouse was set at 100% and decreased performances normalized against this value. For assessment of survival, animals were kept until end-stage. Once the animals got severely impaired hind limbs (typically around ∼P140), they were monitored twice a day and scored using the KI assessment checklist to follow animal health status. From this point water bottles with long drinking spouts were provided to facilitate water access. In addition, animals received soft gel food (Scanbur) on the cage floor. End stage was defined as the inability of the animals to rise within 15 seconds once they lie on both sides.

### Tissue collection, processing and analysis

For muscle analysis, mice were sacrificed by inhalation of CO_2_. Lumbrical muscles (from the plantar surface of the hind-paw), soleus and tibialis anterior (TA) were dissected in 0.1□M phosphate buffered saline (PBS) and fixed in 4% paraformaldehyde (PFA) (Sigma-Aldrich) for 30□minutes for NMJ analysis. Lumbricals were analyzed in whole-mount, whereas soleus and TA muscles were sectioned at 30□μm thickness. For CNS immunohistochemistry, animals were anesthetized with avertin (2,2,2-Tribromoethanol; Sigma-Aldrich) and perfused intracardially with PBS followed by 4% PFA. Brains and spinal cords were dissected and postfixed (for 3□hours and 1□hour, respectively), cryoprotected in sucrose and sectioned (30□μm). Tissues were imaged on a Zeiss LSM700 or 800 confocal microscopes and a Zeiss Axio imager M1 microscope.

### Quantification of GAP-43 expression at the NMJ

For analysis of GAP-43 expression at the NMJ the first and second deep lumbrical muscles from the hind paw of P140 SOD1^G93A^;β2m+/+ and SOD1^G93A^;β2m-/- mice were dissected whole mount to preserve the entire innervation pattern. After fixation, tissue was permeabilized in 4% triton-X 100 (Sigma; 0.1%) in 0.1M PBS for one hour and blocked in 10% donkey serum (Jackson Immuno Research) and 0.1% triton-X 100 in 0.1M PBS for a further hour at room temperature. Muscles were incubated over two nights at 4°C in blocking solution with primary antibodies directed against neurofilament (DSHB, 2H3, 1:50) and GAP-43 (Millipore, AB5520 1:250) in order to visualize axons and regenerating neurons, respectively. Muscles were then washed twice for 30 minutes in 0.1% triton-X 100 and 10% donkey serum in 0.1M PBS and incubated for 2 hours with Alexa Fluor 488 donkey anti-mouse and Alexa Fluor 568 donkey anti-rabbit secondary antibodies in 0.1M PBS (1:500, Life Technologies). Muscles were washed in 0.1M PBS for 30 minutes and then exposed to Alexa Fluor 647 alpha-bungarotoxin (alpha-BTX; 1:1000, Life Technologies) for ten minutes to label post-synaptic acetylcholine receptors. Muscles were then whole-mounted in Mowiol 488 (Sigma-Aldrich) on glass slides and cover-slipped for subsequent imaging. Imaging was performed on laser scanning confocal microscopes (Zeiss LSM700 and LSM800). At least 50 NMJs were individually categorized per muscle per mouse based on the level of GAP-43 expression at each one (minimum of 65 NMJs per muscle). GAP-43 levels were classed as distinct (bright and defined staining overlying the endplate), diffuse (faint and undefined staining, or only partially overlying the endplate) or devoid (no GAP-43 overlying the endplate), as previously described (Allodi et al. 2016). All analyses and quantifications were performed blind to the genetic status of the muscles.

### NMJ innervation in tibialis anterior, lumbricals and soleus muscles

Muscle was permeabilized with 4% triton X-100 for 1 hour. Blocking was performed with 10% donkey serum in 0.1% triton X-100 (Sigma Aldrich) in PBS and the tissue was subsequently incubated with primary antibodies in 10% donkey serum and 0.1% triton X-100 in PBS for 48 hours. Primary antibodies used were mouse anti-synaptic vesicle protein (DSHB, SV2, 1:100) and mouse anti-neurofilament 165 kDa (DSHB, 2H3; 1:50). Next, slides were washed with PBS and incubated with secondary Alexa-488 donkey anti-mouse antibodies in PBS (1:500; Invitrogen) for 3 hours at room temperature. Finally, to visualize endplates, α-bungarotoxin (α-BTX) staining was performed for 15 minutes using tetramethyl-rhodamine isocyanate-conjugated α-BTX (1:1,000; Invitrogen). Slides were coverslipped using Mowiol 488 mounting media (Sigma-Aldrich). Innervation of the NMJ was determined by analysing a minimum of 50 endplates across the muscle. Each endplate within a field of view was categorized as either fully occupied (the presynaptic terminal completely overlies the endplate), partially occupied (the presynaptic terminal partially covers the endplate) or vacant (no presynaptic staining overlies the endplate). All analyses and quantifications were performed blind to the genetic status of the muscles.

### Motor neuron counts in mouse spinal cords

For quantification of spinal MN somas, end-stage SOD1^G93A^;β2m+/+, SOD1^G93A^;β2m+/-, SOD1^G93A^;β2m-/- and age-matched WT;β2m-/- mice (2-4 animals per genotype and sex) were analyzed. Spinal cords were cut at the lumbar level, mounted in OCT and sectioned (30 µm-thickness) in a cryostat. Lumbar sections were attached to Superfrost Plus slides (Thermo Scientific) and stored at −20°C. Prior to staining, slides were air-dried, fixed in 25% ethanol for 2 minutes and Nissl-stained using 2% Cresyl Violet acetate (Sigma-Aldrich, in 25% ethanol, pH8) for 2 minutes. Slides were washed in water for 1 minute and then dehydrated in ethanol 25%, 50%, 70%, 95% and 100% for 2 minutes each. Finally, slides were moved to Xylene for 3 minutes, quickly air-dried, mounted using Mountex (Histolab) and kept at room temperature. MNs were counted in a minimum of six pairs of ventral horns at the lumbar spinal cord. Cells were counted on a Zeiss Axio Imager M1 Upright microscope using Q-capture software. In order to selectively count alpha-MNs, only cells located in the ventral horn with a cell body diameter greater than 20 μm and a clear nucleolus were included. To ensure accurate assessment of cell body size the projected image in Q-capture was calibrated to match a grid rendered on acetate which could be superimposed over the image of the ventral horn so that each box measured precisely 20 μm. This way only cells with a cell body diameter greater than 20 μm were counted.

### Immunohistochemistry in human tissue

Paraffin-embedded human sections of 12 μm were incubated at 60°C for 30 minutes, then moved to xylene for 10 minutes at room temperature. They were sequentially rehydrated with 100% ethanol (2 x 10 min), 95% ethanol (2 x 10 min), 70% ethanol (5 min), 50% ethanol (2 min), 25% ethanol (2 min), and finally rinsed with deionized water for 1 minute. Slides were heated in citrate buffer (10mM Sodium Citrate, Sigma-Aldrich S4641; 0.05% Tween 20, Sigma-Aldrich P7949; pH 6.0) at 95°C for 20 minutes and washed in PBS for 30 minutes. Endogenous peroxidases were quenched using a 50% methanol, 3% hydrogen peroxide solution (H_2_0_2_; Sigma Aldrich, 216763) in PBS for 10 minutes, followed by a 5-minute PBS wash. Sections were blocked with 10% serum in PBS with 0.1% Triton X-100 (Sigma Aldrich) for 1 hour, and then incubated with primary antibody in PBS with 0.1% Triton X-100 and 10% serum for 72 hours at 4°C. Primary antibodies used for immunohistochemistry in human tissue were rabbit anti-GFAP antibody (Dako, Z0334, 1:200) and mouse anti-HLA Class 1 ABC antibody (Clone EMR8-5, Abcam, ab70328, 1:100). For HLA staining, both blocking and primary antibody incubation were carried out without Triton X-100. After washing three times in PBS, slides were incubated with Biotin-SP affinitypure donkey anti-rabbit IgG (Jackson ImmunoResearch, 711.065.152) or anti-mouse IgG secondary antibody (Jackson ImmunoResearch, 715.065.151) in PBS overnight at 4°C. The avidin-biotin complex (ABC) (VECTASTAIN Elite ABC-HRP Kit, Vector Laboratories PK-6100) solution was applied for 1 hour at room temperature, followed by washes in PBS (3 x 10 min). DAB staining was performed using the Vector DAB substrate kit (Vector Laboratories, SK-4100), with a final wash in ddH□O for 10 minutes. Nuclei were counterstained with Myers hematoxylin (Sigma Aldrich, GHS132) for 2 minutes, rinsed in ddH□O, then treated with 70% ethanol containing 36 mM HCl to remove background staining, followed by another ddH□O rinse. Dehydration was completed with sequential ethanol treatments for 2 minutes each (25%, 70%, 75%, 95%, and 100% ethanol), xylene (2 x 2 min), and mounted using DPX new non-aqueous mounting medium (Sigma Aldrich, 1005790500). Images of spinal cord and brain sections were acquired with a Zeiss Axio Observer 7 microscope under bright□field conditions at 20×.

### Protein level analysis in human tissue

The human *post mortem* tissues used for quantification of HLA and GFAP stainings are listed in Supplementary Tables S1-2. For quantification of HLA staining intensity in MNs in spinal cord and midbrain the following procedures were followed. Neurons located in the ventral horn of the spinal cord were manually outlined using the ImageJ software (NIH, Bethesda, MD), and the intensity of the staining was measured in somas with an area of 300□μm^2^ or greater, which were considered to be MNs based on size and location, and the background staining was subsequently subtracted. OMNs were identified based on their anatomical location in midbrain tissue sections and morphological features, manually outlined in ImageJ, and staining intensity in somas was quantified after background subtraction. The absolute majority of OMN somas had an area ranging from 200 μm^2^ to 1200 μm^2^ (Supplementary Figure 1F). The number of donor samples used and MNs quantified for HLA staining are outlined in Supplementary Table S3.

For GFAP staining around spinal MNs, images were taken in the ventral horn of the spinal cord where MNs could be identified, and the mean intensity of each image was measured using ImageJ software, and the background staining subtracted. For GFAP quantification around OMNs, images were taken in midbrain sections, and staining intensity for each image was calculated as the average of the mean intensity from three equal-sized regions of interest. The number of donor tissue samples, and sections used from these as well as images captured and quantified are listed in Supplementary Table S4.

### Statistics

Data was collected and analyzed using GraphPad Prism software. Statistical significance was determined as follows. For the MNs size analyses the Kolmogorov-Smirnov test was performed. For the measurement of staining intensities in MNs the Mann-Whitney Unpaired t-test was performed. For correlations of intensity with MN size, Spearman correlation was performed. For the mRNA levels and NMJ analyses, experimental data were compared by 2-way ANOVA. For behavioral analysis, mice were compared using ANOVA (Kruskal–Wallis test) followed by Dunn–Bonferroni post hoc correction. The survival comparison was performed using Matel-Cox test. For analyses of MN numbers, one□way ANOVA followed by a *post hoc* Tukey was performed.

## Results

### HLA protein is reduced in ALS patient MNs while β2m and HLA mRNAs show compensatory upregulation

We first quantified the number of spinal MNs in *post mortem* tissue from control and ALS donors. We observed a loss of MNs across all sizes. The larger diameter MN somas (>2800μm^2^), which were fewer in number to start with, were completely lost in ALS patients (Figure 1B, *P<0.05*). To understand the role of MHC-I in ALS, we analyzed if there were changes in HLA protein levels in spinal MNs with disease. Quantification of immunostaining demonstrated a significant reduction in HLA protein in remaining spinal MNs of ALS patients compared to controls (Figure 1C-D, *P*<0.0001) and Supplemental Figure 1A-B) which was unrelated to the donor age of the tissue (Supplemental Figure 1C). The decline in HLA levels was apparent across MN sizes between ALS and Control (Figure 1E, *P<0.0001*). Furthermore, there was a clear inverse correlation between HLA protein expression and MN size in both control and ALS tissues (Figure 1F, Ctrl (top panel): Spearman correlation, R=-0.31, *P<0.0001*; ALS (bottom panel): Spearman correlation, R=-0.59, *P<0.0001*).

Next, we evaluated the expression of the astrocytic marker GFAP, which is known to be induced during inflammation in ALS. We observed an increased intensity of GFAP, demonstrating glial activation and neuroinflammation, surrounding spinal MNs in ALS patient tissues (Figure 1G,H, *P<0.0001*, Supplemental Figure 1D,E). These results corroborate and extend upon previous findings showing a reduction in HLA protein levels in spinal motor neurons in end-stage ALS patient tissues, which was shown to render MNs susceptible to activated astrocyte-toxicity (Song et al. 2016).

In light of the fact that MHC-I levels may underlie spinal MN susceptibility in ALS, we investigated the HLA expression in oculomotor neurons (OMNs), which are resistant to degeneration in ALS (Figure 1I,J). Quantification of HLA protein expression and number of OMNs showed no significant difference between control individuals and ALS patients in protein levels, loss of neurons or change in soma sizes (Figure 1I,J; Supplemental Figure 1F,G). This inversion shows that, in healthy tissue, larger OMNs express the highest levels of HLA; however, in ALS, the largest OMNs show the least HLA expression (Supplemental Figure 1H). This indicates that while the overall average expression is maintained (Figure 1 I,J), OMNs are not entirely unaffected in ALS. Furthermore, the observation that there was no increased GFAP immunoreactivity detected immediately around OMNs (Figure 1K,L; Supplemental Figure 1I) suggests that this initial HLA dysregulation might occur before any astrocytic activation and within a non-inflammatory microenvironment. This demonstrates that OMNs persist in ALS, maintain overall normal MHC-I expression and are not exposed to the same inflammatory environment as spinal MNs in ALS, potentially contributing to their resistance to degeneration. The overall maintenance of HLA levels in OMNs, in contrast to their reduction in vulnerable MNs, supports the hypothesis that MHC-I downregulation is selectively associated with neuronal populations undergoing degeneration. To further explore the link between MHC-I signaling and differential susceptibility to ALS, we conducted a thorough investigation of *β2m* and *HLA* expression levels across MN subpopulations that show differential vulnerability to ALS. We first reanalyzed data from several published studies that isolated MNs using laser capture microdissection (LCM) followed by microarray or RNA sequencing in control mouse or human tissues. This analysis showed that *β2m* mRNA expression levels were equal in disease-resistant OMNs and Onuf’s nucleus MNs and vulnerable spinal MNs in mouse ((Kaplan et al. 2014)*, GSE52118*) (Supplemental Figure 2A) and human ((Allodi et al. 2019), GSE93939) (Supplemental Figure 2B) ((Brockington et al. 2013)*, GSE40438*) (Supplemental Figure 2C), and so were *HLA* levels (Supplemental Figure 2A-C). Thus, the basal mRNA expression level of *β2m* and *HLAs* in healthy neurons do not explain differential susceptibility to ALS.

Neurons show both protective and detrimental responses to disease which can explain their early demise or resilience towards ALS (Mei et al. 2025; Nichterwitz et al. 2020; Perrin et al. 2005). It has been reported that *β2m* mRNA levels increase with disease severity in spinal MNs of SOD1^G93A^ mice (Staats et al. 2013). This may at first seem contradictory to the drastic down-regulation of HLA protein in ALS ((Song et al. 2016) and Figure 1D). However, these findings may rather point to compensatory regulation of components of the MHC-I complex in ALS in response to the loss of the respective proteins, that may play a role in ALS disease progression. We thus conducted bioinformatics analyses of a number of published studies comparing control and ALS tissues across MN subpopulations and species. We continued to analyze our own published data sets (Allodi et al. 2019; Nizzardo et al. 2020) where we conducted RNA sequencing on human lumbar spinal MNs and OMNs isolated from *post mortem* tissues from control and ALS donors. In end-stage ALS donor tissues, remaining spinal MNs had a significantly higher level of *β2m* than control MNs (Figure 1M, *P*=0.03, N= 12 (Ctrl), N= 7(ALS)), while the level in OMNs remained unchanged and significantly lower than in ALS spinal MNs (Figure 1M, *P=0.09,* Welch t-test, N=12 (Ctrl), N=4 (ALS)). Thus, maintaining low levels of *β2M* alone does not seem to put MNs at risk. *HLA-A and HLA-C* levels were significantly higher in ALS spinal MNs than in control and *HLA-A* was also induced in OMNs in ALS, while *HLA-B* levels remained unchanged in ALS across MN groups (Figure 1O-P). Collectively, there appears to be a discordance between HLA protein and mRNA regulation in ALS patient tissues, with a compensatory upregulation of certain *HLA* mRNAs both in resilient and vulnerable MNs that is not reflected in the protein level. To better comprehend the temporal regulation during disease we next analyzed a number of transcriptome data sets from SOD1 mouse models. Specifically, a microarray data set on lumbar MNs isolated by LCM at 13 weeks (P90) and 17 weeks (P120) from the SOD1G93A mouse (Perrin et al. 2005), showed an increase in *β2m* levels at an early symptomatic stage of 13 weeks (*P=0.01* Welch t-test, N=3) in ALS, but not at the later symptomatic stage 17 weeks (N=3) compared to wild-type littermates, nonetheless there was a significant increase in ALS from 13 to 17 weeks (*P=0.04* Welch t-test, N=3). Here, the large variability across control samples at 17 weeks may have precluded the comparison with the ALS data of the same stage. *H2-K1* levels remained low at 13 weeks in ALS. Although not statistically significant (P=0.16, Welch t-test, N=3), an increase was observed at 17 weeks in ALS, similar to the human equivalent, *HLA-C*, in end-stage sALS patient MNs (Figure 1M-O and Supplemental Figure 2D). An RNA sequencing study on spinal MNs isolated from presymptomatic SOD1G85R mice at three months of age (shows paralysis at 5-6 months of age) ((Bandyopadhyay et al. 2013)*, GSE38820*) showed an increase in *β2m* levels and *HLA-A* with disease, while H2-K1 remained unchanged (N=2) (Supplemental Figure 2E). A microarray data set on spinal MNs isolated from 15-week-old presymptomatic SOD1G37R mice (shows paralysis at 25 weeks) and wild-type littermate controls (Lobsiger et al. 2007) showed maintained expression levels of *β2m*, *H2-D1* and *H2-K1* in disease (Figure 1N, O and Supplemental Figure 2F). In conclusion, our comprehensive transcriptome analysis across multiple datasets shows that *β2m* and *HLA-C* is consistently upregulated in sALS in spinal MNs, as well as in symptomatic stages of ALS mice (*H2-K1*, mouse equivalent to *HLA-C* in humans), while *HLA-A* was only found upregulated in sALS MNs (Figure 1M-P).

Collectively, our findings show that the largest spinal MNs exhibit the lowest level of HLA protein and this expression is lost in ALS, while the disease-resilient OMNs maintain HLA levels in ALS, is a compelling correlation with selective vulnerability and resilience. Coupled with the apparent compensatory upregulation of *β2m* and *HLA-C* mRNAs in sALS patient MNs and the intriguing differences in outcome of β2m knockout in ALS made us pursue analysis of the effect of β2m loss in ALS.

### Knockout of β2m leads to a small, temporary worsening of muscle strength but has no impact on survival

To investigate the impact of β2m on ALS we crossed β2m knockout (β2m-/-) mice with SOD1G93A mice, both on C57/Bl/6 background (Figure 2A). We analyzed motor performance by grid delay test and extension reflex. We observed a significant worsening in SOD1G93A males with the β2m-/- background from 3 to 5 weeks in the latency to fall after maximum performance (Figure 2B, ANOVA (Kruskal–Wallis test followed by Dunn–Bonferroni post hoc correction); *P*=0.0348 (week 3), *P*=0.00429 (week 4), *P*= 0.0124 (week 5; compared to β2m+/-), N=7-12/group/sex). In females, we observed a significant worsening of maximum performance in SOD1G93A females with the β2m-/- or β2m+/- background after 4 weeks compared to the β2m+/+ background (Figure 2C, ANOVA (Kruskal–Wallis test followed by Dunn–Bonferroni post hoc correction); *P*=0.0278 compared to β2m+/-, *P*= 0.0492 compared to β2m-/-, N=7-12/group/sex). No effect in the extension reflex of either group or sex was found (Figure 2D, E). (N=7-12/group/sex). All the SOD1G93A mice showed a clear weight loss compared to the β2m-/- lacking SOD1G93A expression, as expected (Figure 2F, G). There was no statistical difference between the β2m genotypes on the SOD1^G93A^ background, for males (Figure 2F, N=9-17/group) or females (Figure 2G, N=14-26/group). The loss of β2m did not give rise to any survival effect in male (Figure 2H, N=8-14/group) or female SOD1G93A (Figure 2I, N=12-17/group) mice (Mantel-Cox test). To investigate if end-stage ALS mice show a similar loss of MNs irrespective of the β2m genotype, and thus could be expected to have reached end-stage due to MN disease, MNs were counted based on Nissl staining of spinal cord sections across genotypes. Quantifications showed a clear decrease in MN numbers in SOD1G93A mice with no difference between the β2m genotypes, compared to the β2m mice on a control background (which have a normal life-span) (Supplemental Figure 3A-G, N=4-5 mice (male and female) per group, *P*=*0.0006*, One-way ANOVA). In conclusion, removing β2m from SOD1G93A mice and thus disabling MHC-I expression from all cells did not significantly change the outcome of disease in the mice, but gave a small, transient, functional worsening in males and females.

**Figure 2.**
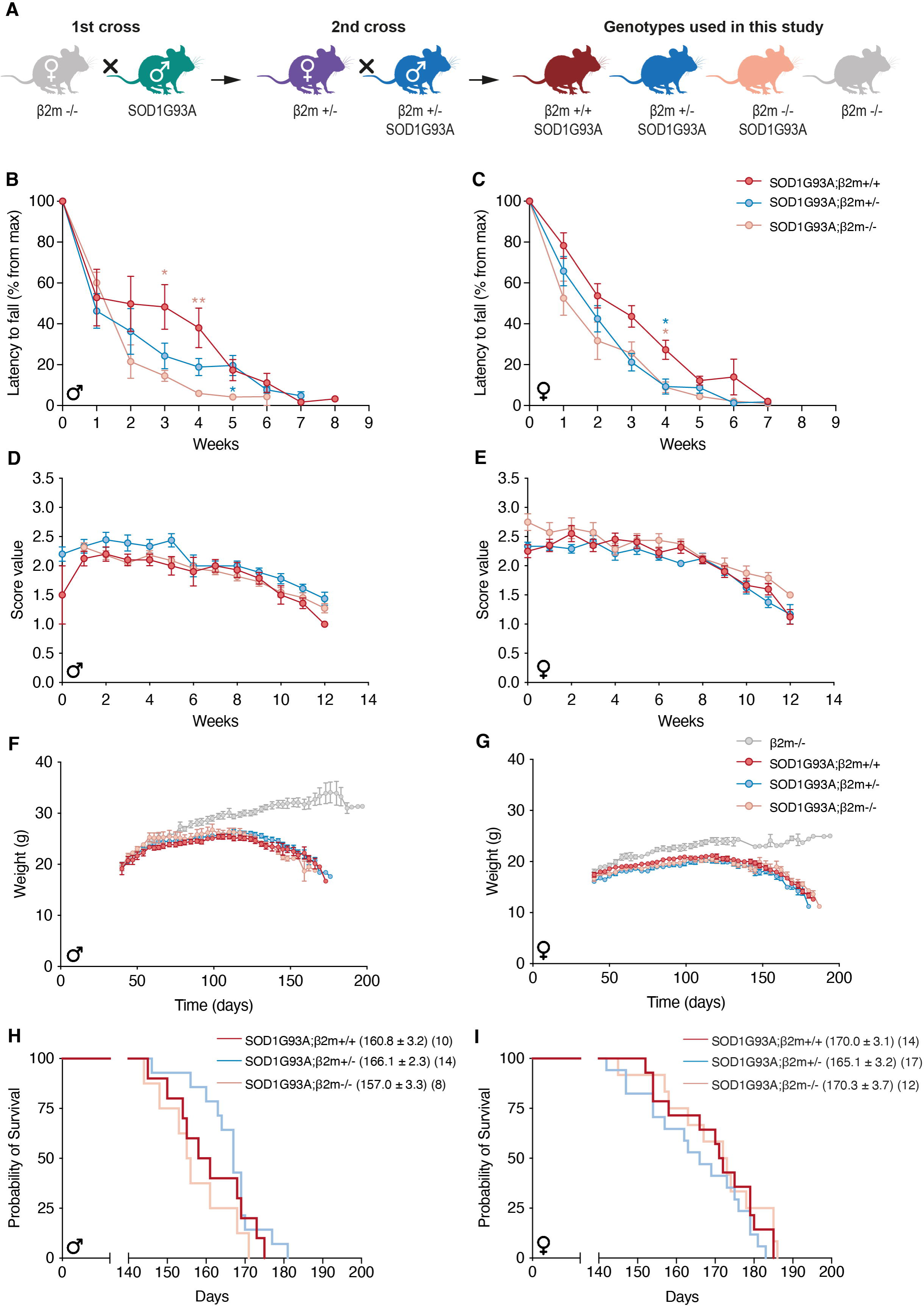
Loss of β2m results in a temporary, early worsening of motor behavior in ALS mice. (**A)** Schematic of mouse crosses to retrieve SOD1G93A mice lacking β2m. Motor performance was assessed by (**B,C**) the latency to fall in the grid test (n=5-6 per group/sex) and (**D,E**) the extension reflex (n=7-12 per group/sex) in male (**B,D**) and female (**C,E)** ALS mice with different doses of β2m. In the grid test (where 0 weeks on the x-axis represents the time point of maximum performance) SOD1G93A males lacking β2m temporarily performed worse (between 3-5 weeks after maximum performance) than SOD1G93A males with normal or heterozygous levels of β2m (ANOVA (Kruskal–Wallis test) followed by Dunn–Bonferroni post hoc correction; *P*=0.0348 (week 3), P=0.00429 (week 4), *P*= 0.0124 (week 5), N=7-12/group/sex; data are expressed as the mean ± SEM). SOD1G93A females lacking one or two alleles of β2m temporarily performed worse at 4 weeks after maximum performance compared to the β2m+/+ background (ANOVA (Kruskal–Wallis test, followed by Dunn–Bonferroni post hoc correction; *P*=0.0278 compared to β2m+/-, *P*= 0.0492 compared to β2m-/-, N=7-12/group/sex; data are expressed as the mean ± SEM). Analysis of extension reflex (where 0 weeks corresponds to 8 weeks of age) showed no change with loss of β2m in either males (**D**) or females (**E**), neither was there an effect on weight loss in either sex (**F-G**). Analysis of survival showed no significant improvement in either sex with knockout of β2m in SOD1G93A mice (**H,I**).

### Knockout of β2m preserves innervation of particular muscles in SOD1G93A ALS mice

To investigate if the removal of β2m in SOD1G93A had any impact on denervation, we conducted an analysis of neuromuscular junctions (NMJs) in tibialis anterior, lumbricals and soleus muscles at P140. The incoming motor nerves were visualized using antibody staining against neurofilament (NEFM) and synaptic vesicle glycoprotein 2A (SV2A), while muscle endplates were labeled with alpha-bungarotoxin (BTX). Representative images of lumbrical muscles from female SOD1G93Axβ2m^+/+^ (Figure 3A, B) and SOD1G93Axβ2m^-/-^ (Figure 3C, D) mice at P140 illustrate these stainings. NMJ innervation patterns were defined as fully occupied, partially occupied or vacant endplates. The innervation in tibialis anterior (Figure 3E) and soleus muscles (Figure 3G) remained unchanged by the removal of β2m. Surprisingly, lumbrical muscles in SOD1G93Axβ2m^-/-^ mice retained their innervation better than in SOD1G93Axβ2m^+/+^, as shown by their higher level of full NMJ occupancy and lower percentage of vacant endplates (Figure 3F, N= 4-5 female mice/group, *P<0.05*, ANOVA). As these mice performed slightly worse in the grid delay test than the mice with the β2m wild-type background it is evident that this innervation did not give a functional outcome in the test utilized. To determine if the higher level of occupancy of NMJs in lumbricals was due to preservation or rather regeneration and subsequent reinnervation of vacated endplates, we stained muscles with an antibody against GAP43 and endplates were again visualized by BTX. This analysis demonstrated that regeneration was similar in either genotype (Figure 3H-J) and thus the higher level of occupied endplates in the SOD1G93Axβ2m^-/-^ is due to increased stability. In conclusion, HLA and β2m are clearly regulated in vulnerable MNs in ALS, with loss of HLA protein and compensatory mRNA upregulation. Removal of β2m results in site-specific preservation of innervation, but is not sufficient to impact motor behavior or survival in ALS mice.

**Figure 3.**
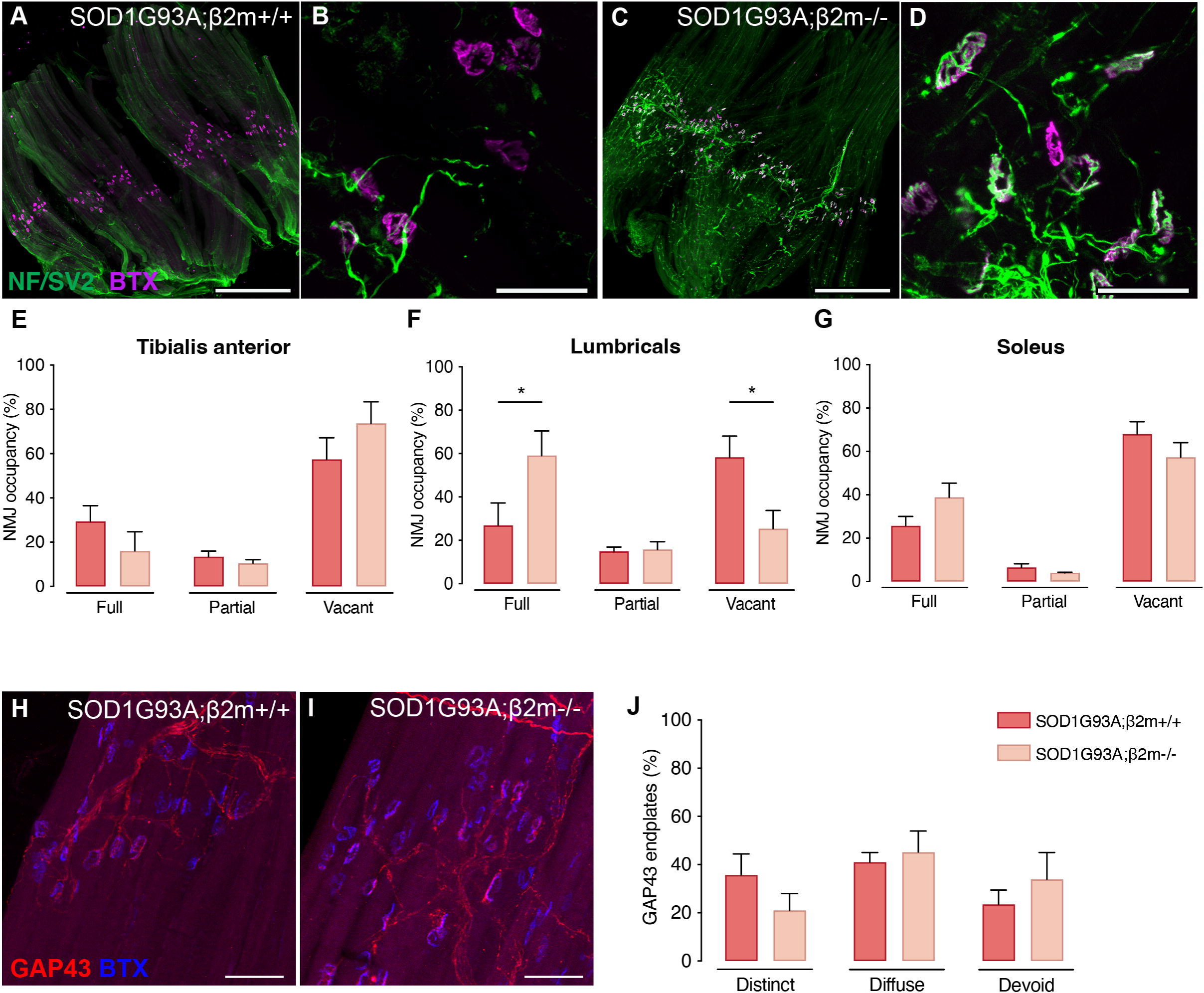
Knockout of β2m preserves innervation of particular muscles in SOD1G93A ALS mice. Representative images of the innervation pattern of motor nerves in lumbrical muscles of female (**A, B**) SOD1G93A;β2m^+/+^ or (**C, D**) SOD1G93A;β2m^-/-^ mice at P140. The incoming motor nerves are visualized by a combined antibody staining against Neurofilament (2H3) and SV2A (green) and muscle endplates are stained with alpha bungarotoxin (magenta). Quantification of innervation patterns, into fully occupied, partially occupied or vacant endplates was conducted in (**E**) tibialis anterior, (**F**) lumbricals and (**G**) soleus muscles. Lumbrical muscles in SOD1G93A;β2m^-/-^ mice retained their innervation better than in SOD1G93A;β2m^+/+^, as shown by (**F**) their higher level of full NMJ occupancy and lower percentage of vacant endplates (N=4-5 female mice/group, *P<0.05*, ANOVA; data are expressed as the mean ± SEM). Representative immunofluorescence images of lumbricals muscles from P140 female (**H**) SOD1G93A;β2m^+/+^ or (**I**) SOD1G93A;β2m^-/-^ mice stained with the regeneration marker GAP43 (red). Muscle endplates were stained with alpha bungarotoxin (blue). (**J**) Analysis of GAP43 innervation to muscle end plates showed no statistical significance between β2m genotypes in any of the innervation pattern classification (N=4 mice/group. Statistical test: 2-way ANOVA; data are expressed as the mean ± SEM). Scale bars: 500 μm (in A, C and H) and 50 μm (in B, D and I).

## Discussion

The selective vulnerability of MNs is a hallmark of ALS, yet the underlying mechanisms remain unclear. While the large and fast-fatiguing MNs are selectively lost early in the disease, resistance is observed among MNs in the oculomotor and Onuf’s nuclei (Comley et al. 2015; Nijssen et al. 2017; Frey et al. 2000). Our study investigated the role of β2m and its associated complex, MHC-I, in dictating this selective vulnerability, building upon complex and often contradictory implications in the existing literature. We first validated the loss of the largest spinal MNs in end-stage ALS, and the persistence of OMNs. A key mechanistic insight from our study focuses on the differential basal expression of MHC-I in MN subpopulations. By examining control donor tissue, we demonstrated an inverse correlation between HLA protein level and MN size in the spinal cord, where the large, most vulnerable spinal MNs exhibited the lowest HLA expression. Beyond ALS, MHC-I dysregulation has been implicated in several neurodegenerative disorders. In Alzheimer’s disease, MHC-I destabilization by amyloid-β leads to synaptic dysfunction (Kim et al. 2023). In Parkinson’s disease, upregulated neuronal MHC-I provokes autoimmune activation, rendering dopamine neurons more susceptible to neurodegeneration (Wang et al. 2021).

Staats et al. demonstrated that β2m expression was upregulated in MNs with disease progression in SOD1G93A ALS mice and that global deletion of β2m shortened survival. This led to the conclusion that β2m may play a protective role in ALS. The authors hypothesized that β2m promotes plasticity critical for muscle reinnervation, a compensatory mechanism that, when lost, accelerates muscle denervation and disease severity (Staats et al. 2013). Song and colleagues on the other hand, demonstrated that MHC-I was downregulated on MNs in symptomatic SOD1G93A mice and on spinal MNs in end-stage ALS patient tissues (Song et al. 2016). As β2m levels were not analyzed by *Song et al*. and HLA levels were not investigated by Staats and colleagues, these findings are not contradictory per se, but may just indicate that while HLA levels are reduced there is compensation on the mRNA level by β2m to try to restore MHC levels.

Our broader transcriptome analysis across data sets indeed showed that both *β2m* and *HLA* mRNAs show compensatory upregulation in ALS, when HLA protein is gradually lost. Reduced or low MHC-I levels may impair homeostatic or regenerative responses, aligning with its emerging non-immune functions in synaptic plasticity and axonal stability (Boulanger 2004; Shatz 2009) and render neurons more susceptible to inflammatory or excitotoxic insults.

In line with the findings of Song and colleagues, we show that HLA protein is markedly reduced in vulnerable spinal MNs in end-stage ALS patient tissues, while we demonstrate that OMNs retained HLA protein levels.

Collectively, these findings strongly indicate that the selective vulnerability of spinal MNs in ALS is not dictated by differences in basal gene transcription of *β2m* or *HLA*, as we found that the mRNA levels of these genes were equal across vulnerable and resilient MNs in control donor tissue. Furthermore, in ALS donor tissue, our observation that spinal MNs upregulate β2m and HLA-C mRNAs, while HLA protein levels were decreased, suggests that post-transcriptional, translational, or localization mechanisms prohibit an increase of HLA protein levels, but this remains to be further investigated in detail. It also shows that vulnerable MNs develop an intrinsic, yet ultimately unsuccessful, protective response, which likely reflects a neuronal stress response, similar to what occurs after axonal injury (Lindå et al. 1998; Thams et al. 2009). This aligns with comparative transcriptome data showing that vulnerable MNs in ALS react with both protective and degenerative gene programs. This was demonstrated by comparing the transcriptome data from Mei et al. (2025) and Shadrach et al. (2021), where the vulnerable MNs shared regeneration-associated factors, such as *Sox11, Gap43, Sprr1a, Adcyap, Chl1, Atf3,* and *Tubb3*, with MNs undergoing axonal repair after crush injury. This suggests that the upregulation of β2m and MHC-I genes could be part of a broader, but insufficient, intrinsic program initiated by the vulnerable MNs to promote plasticity and survival.

Song et al. demonstrated that ALS astrocytes reduced MHC-I expression on MNs, rendering these more vulnerable to astrocyte-mediated toxicity. Conversely, increasing MHC-I expression on MNs, for instance by overexpressing the human MHC-I molecule HLA-F, increased survival and improved motor performance *in vivo*. This established a paradigm where the maintenance of MHC-I expression is an active determinant of MN survival in the toxic ALS microenvironment. In line with these findings of an interplay between cell autonomous and non-cell autonomous mechanisms regulating vulnerability in ALS, we found no increase in astrocytic GFAP immunoreactivity around OMNs in ALS while the correlation between HLA protein levels and OMN size shifts from positive in control donors to inverse in ALS. This supports the idea that the preservation of a healthy glial milieu contributes to OMN resistance. These observations imply that in vulnerable spinal MNs, an early dysregulation of HLA expression may act as a trigger for pathological neuron-glia crosstalk, leading to glial activation and, ultimately, neurodegeneration. In contrast, the maintenance of overall HLA expression and the absence of reactive gliosis in OMNs likely protect these neurons from entering the same degenerative cascade. This aligns with the concept that non–cell-autonomous mechanisms, particularly astrocyte-induced toxicity, contribute to selective MN loss, and that their activation is induced by cell autonomous mechanisms in vulnerable neurons (Clement et al. 2003; Boillée et al. 2006; Yamanaka et al. 2008; Ilieva et al. 2009). The differential loss of MHC-I protein in vulnerable and resistant MNs could be directly linked to these differences in local microenvironment, as Song et al. established that toxic ALS astrocytes reduce MHC-I expression on MNs, making these susceptible to astrocyte-induced cell death. Our finding that the largest, most vulnerable spinal MNs show the lowest HLA protein levels, combined with the neurotoxic environment created by activated spinal astrocytes, indicate that this puts them at a particular risk in ALS.

Finally, Nardo et al. (2018) highlighted that MNs are particularly responsive in upregulating MHC-I molecules in response to insults, and this response is associated with preservation of axonal structure and function and slower disease progression in ALS mice. They demonstrate that MHC-I depletion accelerated disease onset and motor deficits but increases overall survival and promotes MN survival in SOD1G93A mice suggesting that MHC-I plays a deleterious role in disease survival. This depletion has different effects on the innervation depending on the targeted muscle groups.

Our current study introduces nuances to this paradigm, suggesting that the beneficial effects of β2m/MHC-I may be more transient and localized than previously assumed, and provides insight into their role in selective vulnerability. Despite the role suggested by these molecular findings, functional genetic ablation of β2m in SOD1G93A mice produced only a transient, modest impairment in motor performance without altering overall survival or MN loss. Additionally, some muscles, such as lumbrical muscles, did exhibit site-specific preservation of innervation, which underscores the muscle- and context-dependence of MHC-I. The preservation observed in fast-fatigable muscles like the lumbricals (typically among the first to degenerate) suggests that loss of β2m may indirectly stabilize NMJs in select circuits. This site-specific activity of MHC-I is reinforced by Nardo et al. where MHC-I depletion accelerated or delayed denervation depending on the targeted muscle groups. Taken together, these results support a model where MHC-I components, including β2m, are dynamically regulated in ALS, exerting a context-dependent role that varies across MN subtypes, disease stages, and peripheral targets.

While our data support a role for MHC-I in selective vulnerability, the functional impact of experimentally manipulating β2m in vivo has been strikingly inconsistent across studies. Four different studies, including ours (Staats et al. 2013; Song et al. 2016; Nardo et al. 2018) have evaluated the impact of removing β2m on disease progression in the SOD1G93A mouse model, resulting in distinct outcomes. Specifically, two studies reported a worsened phenotype and shortened survival (Staats et al. 2013; Song et al. 2016); one reported an improved phenotype and increased survival (Nardo et al. 2018); and our study found no significant impact on overall survival or MN loss, only a transient functional deficit. In those studies, both the SOD1G93A and the β2m knockout mice were reported to be on a C57BL/6 background. The striking message from this comparison is that the C57BL/6 substrains and/or non-genetic variables appear to have a greater impact on the disease phenotype. It is well-documented that the longevity of mouse strains, even within the C57BL/6 lineage, can lead to phenotypic differences. As highlighted by The Jackson Laboratory (JAX), the longer time that strains are separated from each other, the greater the number of genetic differences between them, which can lead to large phenotypic differences (JAX, 2016, « There Is No Such Thing as a C57BL/6 Mouse! »). In their seminal work, Nardo et al., 2013 (Nardo et al. 2013), demonstrated that SOD1G93A mice on distinct genetic backgrounds (C57BL/6 vs. 129Sv) show consistent differences in disease progression and lifespan. Crucially, the slower-progressing C57BL/6 strain exhibited a striking upregulation of immune system processes and increased MHC-I expression in MNs at disease onset, contrasting with the rapidly progressing 129Sv strain. This establishes that the genetic background modulates key pathways which may then override the subtle experimental effect of the β2m knockout in the different studies. Furthermore, differences in microbiota, environmental stressors, or facility-specific pathogens could further modulate the immune-neuronal interplay and thereby the β2m phenotype. Those could influence the outcome, as the animal’s immune status is impacted by the loss of β2m, a key variable not investigated in any of the studies. Lastly, the high degree of genetic drift among B6 substrains is known to influence neuroinflammatory and metabolic traits (JAX Blog, 2016), reinforcing the conclusion that the background of the animal and immune status likely mattered more than the loss of β2m itself.

Taken together, our findings provide new insights into the complex, context-dependent regulation of MHC-I in ALS. We demonstrate that the HLA protein levels are determinant of MN vulnerability, with the most susceptible MNs exhibiting the lowest HLA protein levels. The observed β2m and HLA-C mRNA upregulation in vulnerable spinal MNs likely represents a compensatory mechanism that, despite the concurrent activation of regenerative factors (Mei et al. 2025), is ultimately overcome by the disease. Our functional in vivo study on SOD1G93A ALS mice lacking β2m suggests that β2m removal does not have a profound, singular effect on disease outcome. While removal of β2m resulted in a modest worsening of functional motor performance, histological analysis suggests a modest functional benefit in NMJ stability and partial preservation of innervation in particular muscles. This indicates the intrinsic protective mechanism mediated by β2m is insufficient to reverse the fatal trajectory of ALS. It is plausible that while MHC-I can protect the distal axon, a function aligned with its role in synaptic plasticity and nerve regeneration (Nardo et al. 2018), this localized protection is not sufficient to counteract the chronic, progressive, and non-cell autonomous toxicity exerted by surrounding glia and mutant proteins (Song et al. 2016). Critically, our comparison with existing literature underscores the overriding influence of genetic and environmental context on ALS phenotypes in mouse models, as demonstrated by the variable effects of β2m removal, likely depending on genetic background and environment.

## Supporting information

Supplementary material, including Supplemental Figure 1-3, and Supplemental Tables S1-4

## Acknowledgements

We would like to thank all members of the Hedlund and Bartosovic laboratories for constructive discussions and helpful suggestions during joint lab meetings. We also want to thank Mattias Karlén for his excellent work on the schematic images in Figures 1A and 2A. Computation and a data handling was enabled by resources provided by the National Academic Infrastructure for Supercomputing in Sweden (NAISS) and the Swedish National Infrastructure for Computing (SNIC) at UPPMAX. This work was supported by grants from the Swedish Research Council and Birgit Backmark’s donation to ALS research at Karolinska Institutet.

