## Supplementary material, including Supplemental Figure 1-3, and Supplemental Tables S1-4 for "Reevaluating the Role of Beta2-Microglobulin: New Insights on Selective Vulnerability in ALS Pathology"

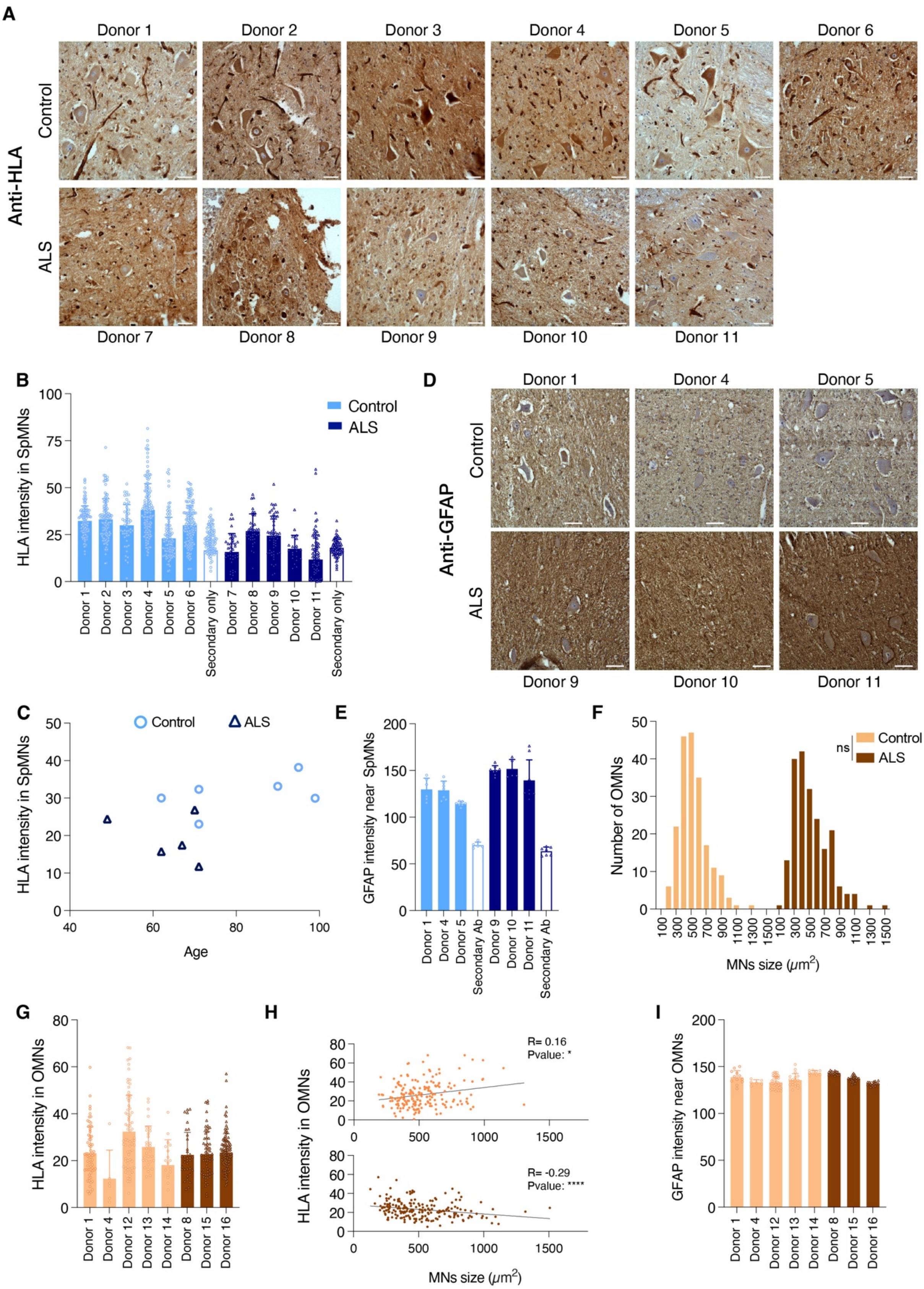
A
Donor 1
Donor 2
Donor 3
Donor 4
Donor 5
Donor 6
Anti-HLA
Control
ALS
Donor 7
Donor 8
Donor 9
Donor 10
Donor 11
B
HLA intensity in SpMNs
Control
ALS
Secondary only
D
Anti-GFAP
Donor 1
Donor 4
Donor 5
Donor 9
Donor 10
Donor 11
C
Control
ALS
HLA intensity in SpMNs
Age
E
GFAP intensity near SpMNs
Secondary Ab
F
Number of OMNs
ns
MNs size ($\mu m^2$)
G
HLA intensity in OMNs
Donor 12
Donor 13
Donor 14
Donor 15
Donor 16
H
HLA intensity in OMNs
R= 0.16
Pvalue: *
R= -0.29
Pvalue: ****
MNs size ($\mu m^2$)
I
GFAP intensity near OMNs

**Supplemental Figure 1. HLA protein level is decreased in MNs in ALS patient tissues while GFAP intensity surrounding MNs is increased.** (**A**) Immunohistochemical staining against HLA-ABC in human *post mortem* spinal cords from control and ALS donors. (**B**) HLA MN intensity across spinal MNs (SpMNs) in individual donor tissues shows a decrease in HLA protein levels in ALS compared to control. (**C**) HLA SpMN intensity shows no correlation with tissue donor age. (**D**) Staining against GFAP to visualize reactive astrocytes in three control and three ALS donor spinal cord *post mortem* tissues (**E**) Quantification of GFAP intensity across SpMNs in individual control and ALS donor spinal cord tissues shows an increase in GFAP levels with ALS, confirming the innate immune reaction. (**F**) There was no significant loss of OMNs in ALS patient tissues or change in soma sizes. (P=0.9993, Kolmogorov-Smirnov test) (**G**) Quantification of HLA protein expression across OMNs in individual donors remained unchanged with ALS. (**H)** HLA immunoreactivity and OMN size are positively correlated in control *post mortem* donor tissues but inversely correlated in ALS tissues. (Ctrl: R=0.16, P=0.0210, Spearman correlation, ALS: R=-0.29, P<0.0001, Spearman correlation) (**I**) There was no increase in GFAP intensity around OMNs in end-stage ALS patient tissues. Data are expressed as the mean ± SD.

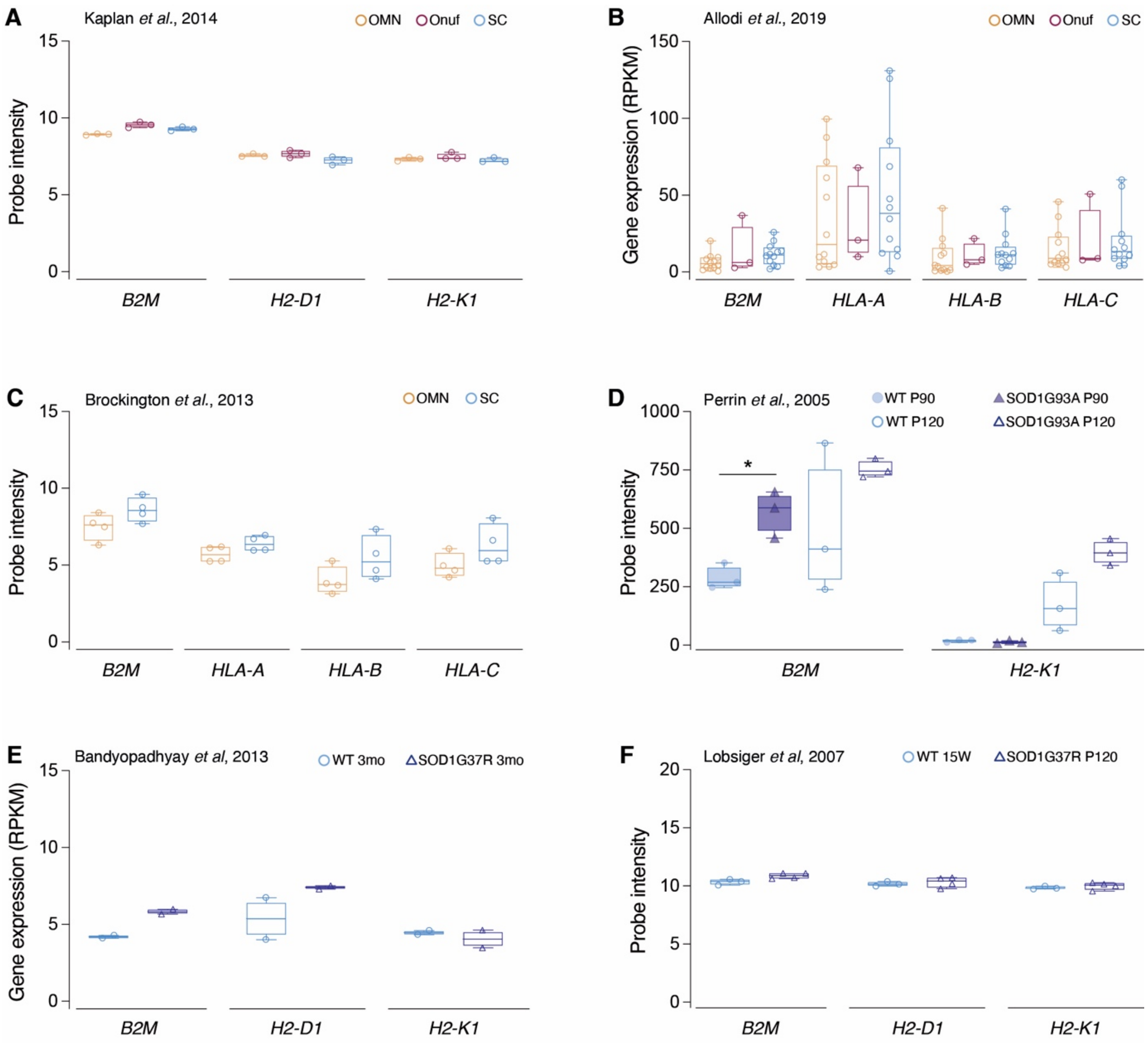


**Supplemental Figure 2. β2m and HLA levels do not underlie differential neuronal vulnerability across brain stem and spinal cord motor nuclei, but are regulated in vulnerable neurons in response to increased disease burden.** Analysis of *β2m*, and *HLA* mRNA levels across control data sets show no statistically significant difference across motor neuron subpopulation with differential vulnerabilities to degeneration in ALS in (**A**) mouse (Kaplan et al. 2014) or (**B, C**) human (Allodi et al. 2019; Brockington et al. 2013). Overexpression of mutant SOD1 in mice causes differential upregulation of β2m and HLA, which may depend on disease stage and mutation, as shown in (**D**) SOD1G93A mice at presymptomatic (P90) and symptomatic (P120) stages (Perrin et al. 2005) and (**E**) SOD1G85R mice at 3 months (Bandyopadhyay et al. 2013) and (**F**) presymptomatic SOD1G37R mice at

15 weeks (Lobsiger et al. 2007). Data are expressed as the median ± min and max. Whiskers extend to 1.5x the interquartile range (IQR).

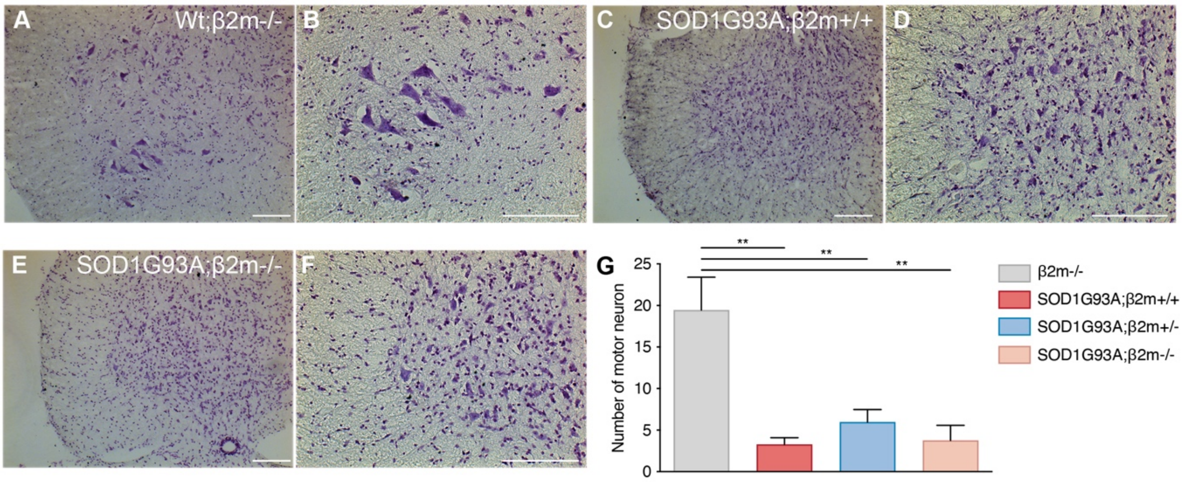


**Supplemental Figure 3. Loss of β2m does not impact the level of motor neuron loss in SOD1G93A ALS mice.** Quantification of spinal motor neurons based on Nissl staining of spinal cord sections across genotypes, with representative bright field images from (**A, B**) β2m$^{-/-}$ mice, (**C, D**) SOD1G93A;β2m$^{+/+}$ mice and (**E, F**) SOD1G93A;β2m$^{-/-}$ mice. (**G**) Summary graph of motor neuron counts showing significance between β2m$^{-/-}$ mice on a wild-type (C57Bl/6) background (*P=0.0006*, one-way ANOVA; data are expressed as the mean ± SEM) and SOD1G93A ALS mice, while no difference was observed between the β2m genotypes in the SOD1G93A ALS mice. Quantifications were done at end-stage for SOD1G93A mice across β2m genotypes and at a matched time point for β2m mice on a control background, which have normal life-span. N=4-5 mice per group (combined male and females).

**Supplemental Table S1.** Human post mortem spinal cord tissue samples used for quantification of anti-HLA and anti-GFAP stainings in and around spinal MNs (relating to Supplementary Figure S1).

| | Spinal cord tissue (SpMNs) | | |
|---|---|---|---|
| | Group | Sex | Age |
| Donor 1 | Control | M | 71 |
| Donor 2 | Control | F | 90 |
| Donor 3 | Control | M | 62 |
| Donor 4 | Control | F | 95 |
| Donor 5 | Control | F | 71 |
| Donor 6 | Control | F | 99 |
| Donor 7 | ALS | M | 62 |
| Donor 8 | ALS | M | 70 |
| Donor 9 | ALS | F | 49 |
| Donor 10 | ALS | M | 67 |
| Donor 11 | ALS | M | 71 |

**Supplemental Table S2.** Human post mortem midbrain tissue samples used for quantification anti-HLA and anti-GFAP stainings in and around OMNs (relating to Supplementary Figure S1).

| | Midbrain tissue (OMNs) | | |
|---|---|---|---|
| | Group | Sex | Age |
| Donor 12 | Control | F | 71 |
| Donor 13 | Control | F | 87 |
| Donor 4 | Control | F | 95 |
| Donor 1 | Control | M | 71 |
| Donor 14 | Control | M | 88 |
| Donor 8 | ALS | M | 70 |
| Donor 15 | ALS | M | 33 |
| Donor 16 | ALS | F | 68 |

**Supplemental Table S3.** The number of human post mortem tissue samples used and MNs quantified for anti-HLA intensity (relating to Supplementary Figure S1).

| | Spinal MNs | | OMNs | |
|---|---|---|---|---|
| | Control | ALS | Control | ALS |
| Number of donors | 6 | 5 | 5 | 3 |
| Number of sections | 24 | 15 | 11 | 6 |
| Number of cells | 550 | 198 | 198 | 158 |

**Supplemental Table S4.** The number of human post mortem tissue samples and sections as well as images captured that were used to quantify anti-GFAP intensity (relating to Supplementary Figure S1).

| | Spinal MNs | | OMNs | |
|---|---|---|---|---|
| | Control | ALS | Control | ALS |
| Number of donors | 3 | 3 | 5 | 3 |
| Number of sections | 15 | 12 | 11 | 6 |
| Number of images | 20 | 21 | 28 | 19 |
